# Linking Branch Lengths Across Loci Provides the Best Fit for Phylogenetic Inference

**DOI:** 10.1101/467449

**Authors:** David A. Duchêne, K. Jun Tong, Charles S. P. Foster, Sebastián Duchêne, Robert Lanfear, Simon Y. W. Ho

## Abstract

Evolution leaves heterogeneous patterns of nucleotide variation across the genome, with different loci subject to varying degrees of mutation, selection, and drift. Appropriately modelling this heterogeneity is important for reliable phylogenetic inference. One modelling approach in statistical phylogenetics is to apply independent models of molecular evolution to different groups of sites, where the groups are usually defined by locus, codon position, or combinations of the two. The potential impacts of partitioning data for the assignment of substitution models are well appreciated. Meanwhile, the treatment of branch lengths has received far less attention. In this study, we examined the effects of linking and unlinking branch-length parameters across loci. By analysing a range of empirical data sets, we find that the best-fitting model for phylogenetic inference is consistently one in which branch lengths are proportionally linked: gene trees have the same pattern of branch-length variation, but with varying absolute tree lengths. This model provided a substantially better fit than those that either assumed identical branch lengths across gene trees or that allowed each gene tree to have its own distinct set of branch lengths. Using simulations, we show that the fit of the three different models of branch lengths varies with the length of the sequence alignment and with the number of taxa in the data set. Our findings suggest that a model with proportionally linked branch lengths across loci is likely to provide the best fit under the conditions that are most commonly seen in practice. In future work, improvements in fit might be afforded by models with levels of complexity intermediate to proportional and free branch lengths. The results of our study have implications for model selection, computational efficiency, and experimental design in phylogenomics.

---

Molecular evolution is heterogeneous across the genome. This poses a challenge for statistical phylogenetic analyses of multilocus data sets, because they rely on explicit models of the evolutionary process (Sullivan and Joyce 2005). There has been considerable interest in the impact of model choice on estimates of evolutionary parameters, such as the tree topology and branch lengths (Steel 2005). For example, an important step in most phylogenetic analyses is choosing a substitution model that captures sufficient variation in the evolutionary process without overfitting the data (Sullivan and Joyce 2005). The task of selecting an appropriate phylogenetic model is especially complex for genome-scale data sets, because the number of potential model combinations becomes astronomical (Lanfear et al. 2012). Therefore, it would be highly beneficial to identify any general principles that can help to improve model fit and performance, while maintaining the tractability of computational analysis.

In terms of model selection in phylogenetics, the models of nucleotide and amino acid substitution have received the largest amount of attention. Various methods have been proposed for identifying the best-fitting partitioning scheme for assigning substitution models to the different loci in the data set (e.g., Lanfear et al. 2012; Kalyaanamoorthy et al. 2017). One aspect of this process that is often overlooked, however, is deciding how to model variation in the pattern of branch lengths of the gene trees. These heterogeneities need to be considered carefully when comparing data-partitioning schemes for phylogenetic analysis. In our descriptions below, we assume that each locus is associated with a gene tree. We also assume that the topologies of these gene trees are identical across loci, such that they can only vary in their absolute length and the pattern of lengths of branches.

The simplest model of branch lengths assumes that they are *universally shared* across loci (Fig. 1a). This model has a length parameter for each of the 2*n*-3 branches in the (unrooted) tree, where *n* is the number of taxa. However, the model is unlikely to be realistic because it assumes that all loci have evolved at identical rates, which contradicts the overwhelming evidence of rate variation across the genome (Bromham and Penny 2003). Nonetheless, this is a widely used model of branch-length variation in molecular phylogenetics. We can generalize the model slightly by allowing loci to have *proportionally linked* branch lengths. In such a model, the branch lengths share proportionality across gene trees, with variation in the summed lengths of these gene trees permitted (Fig. 1b; Yang 1996; Nylander et al. 2004). In other words, all of the gene trees share the same relative branch lengths, but have evolved at different absolute rates. For an unrooted tree, this model of branch lengths has (*L*-1)+(2*n*-3) parameters, comprising a set of 2*n*-3 branch lengths for an arbitrarily chosen gene tree and the *L*-1 relative rates of the remainder of the *L* loci. For each gene tree, the branch lengths can be obtained by multiplying the 2*n*-3 branch lengths of the ‘reference’ gene tree by the relative rate at the locus in question. This pattern in branch lengths can be regarded as the additive outcome of lineage effects and gene effects (Gillespie 1991; Muse and Gaut 1997).

**Figure 1.**
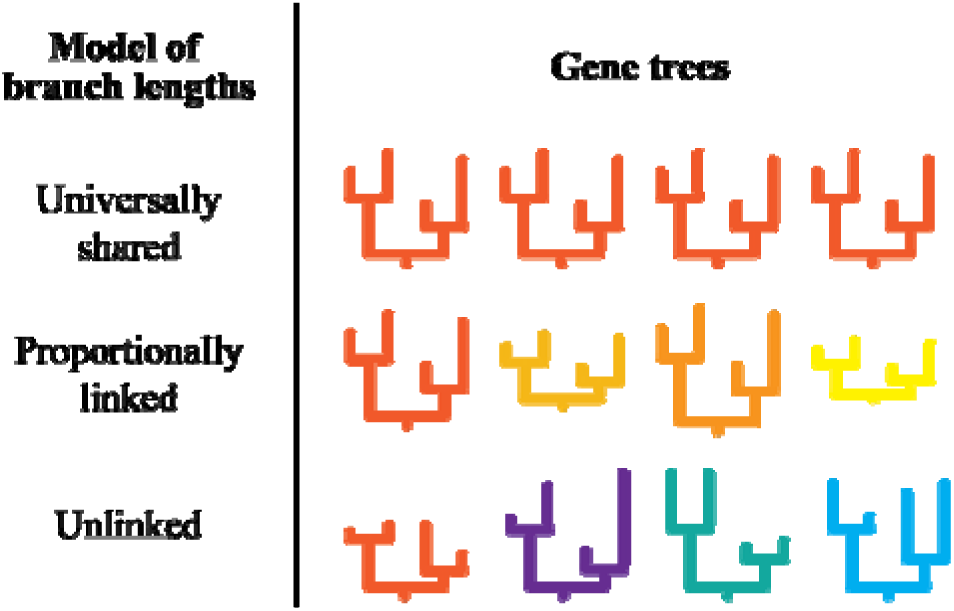
Models of branch lengths across gene trees. A model with *universally shared* branch lengths assumes a single set of branch lengths across gene trees. This model has 2*n*-3 branch-length parameters, where *n* is the number of taxa. A model with *proportionally linked* branch lengths assumes that the proportionality of branch lengths is maintained across gene trees. Nonetheless, variation in the summed branch lengths (tree lengths) is permitted through a scaling parameter per gene tree. This model contains (*L*-1)+(2*n*-3) parameters, where *L* is the number of loci (assuming one gene tree per locus). A model with *unlinked* branch lengths assumes an independent set of branch lengths per gene tree, so it has L·(2*n*-3) parameters.

The third and most parameter-rich model of branch lengths allows each gene tree to have a distinct set of branch lengths (Fig. 1c). This model assumes *unlinked* branch lengths and has *L*·(2*n*-3) parameters. At first glance, this might seem to be the most realistic of the three models of branch lengths because we would expect different loci to evolve under varying degrees of selection and thus to have differing patterns of evolutionary rates across branches (Takahata 1987; Cutler 2000; Ho 2014). However, the number of parameters in the model increases rapidly with the number of loci, meaning that the model will have many parameters when applied to large, multilocus data sets. A biological mechanism that could give rise to this pattern is that in which selective constraints vary among genes and among lineages, known as gene-by-lineage interactions (Gillespie 1991; Muse and Gaut 1997).

The choice of branch-length model has the potential to affect the quality of phylogenetic inference (Marshall et al. 2006). However, the biological basis for choosing among the three models is not well understood. Some studies have suggested that loci vary little in terms of the patterns of branch lengths of their gene trees (Snir et al. 2012, 2014), but others have found evidence of substantial disparities (Bedford and Hartl 2008; Duchêne and Ho 2015).

Here, we compare the statistical fit and performance of the three models of branch lengths in phylogenetic analyses of multilocus data sets. These models vary in terms of whether branch lengths are universally shared, proportionally linked, or unlinked across loci. We combine these models of branch lengths with different partitioning schemes for substitution models. Our analyses of eight multilocus data sets and two phylogenomic data sets show that the best fit is usually provided by a model with proportionally linked branch lengths across loci. We also present a simulation study in which we demonstrate that the fit of the three models of branch lengths depends on the size of the data set.

## Materials and Methods

### Phylogenetic Models Used for Analysis

We analysed a range of multilocus data sets using seven different partitioning treatments for branch lengths and substitution models (Table 1). Branch lengths were assumed to be universally shared (treatments 1–3), proportionally linked (treatments 4 and 5), or unlinked (treatments 6 and 7) across loci. For each model of branch lengths, we considered three methods of partitioning the data and selected substitution models from 88 possible models in the GTR+I+Γ+F family of models specified by the command –m TEST in the IQ-TREE software (Nguyen et al. 2015). First, we assumed a simple model in which all loci shared the same substitution model parameters and parameter values (treatment 1). Second, we used an automatic likelihood-based merging approach to select the partitioning scheme (treatments 2, 4, and 6 in Table 1; Lanfear et al. 2012; Kalyaanamoorthy et al. 2017). Third, we applied a partitioning scheme in which each locus has an independent substitution model (treatments 3, 5, and 7 in Table 1).

**Table 1.** Models of branch lengths across gene trees, compared using multilocus and phylogenomic data sets.

| Treatment number IQ-TREE command | Model of branch lengths | Number of substitution models across loci | Number of tree lengths across loci | Number of branch-length patterns across loci | Potential equivalence to other models |
| --- | --- | --- | --- | --- | --- |
| (1) Not applicable | Universally shared | 1 | 1 | 1 | – |
| (2) -TESTMERGE -q | Universally shared | $1 \leq x \leq L$ | 1 | 1 | 1, 3 |
| (3) -TEST -q | Universally shared | $L$ | 1 | 1 | – |
| (4) -TESTMERGE -spp | Proportionally linked | $1 \leq x \leq L$ | $1 \leq x \leq L$ | 1 | 1, 2, 5 |
| (5) -TEST -spp | Proportionally linked | $L$ | $L$ | 1 | – |
| (6) -TESTMERGE -sp | Unlinked | $1 \leq x \leq L$ | $1 \leq x \leq L$ | $1 \leq x \leq L$ | 1, 2, 4, 7 |
| (7) -TEST -sp | Unlinked | $L$ | $L$ | $L$ | – |

In the treatments with automated model selection, the chosen partitioning scheme had the potential to match those in some of the other treatments. This could occur if the method selected the simplest model, in which all loci shared the same substitution model and the same set of branch lengths. On the other hand, the method could select the most complex model, in which each locus had its own substitution model and own set of branch lengths.

### Multilocus and Phylogenomic Data

We applied each of the seven treatments of branch lengths and substitution models to eight multilocus data sets that represented a diverse range of animals and plants. The data sets were taken from an existing curated compilation of data (Table 2; Kainer and Lanfear 2015), and each comprised nucleotide sequences from between two and six loci. The sequence alignments are available from Figshare (doi.org/10.6084/m9.figshare.991367).

**Table 2.**
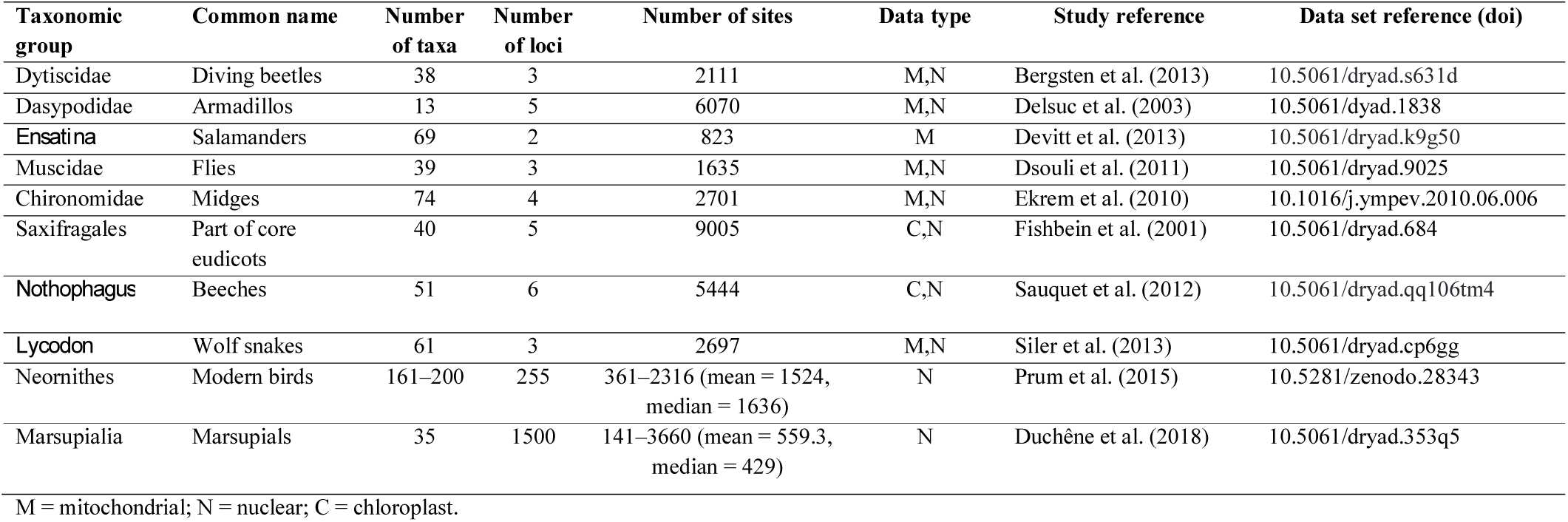
Data sets used for examining models of branch lengths across loci.

| **Taxonomic group** | **Common name** | **Number of taxa** | **Number of loci** | **Number of sites** | **Data type** | **Study reference** | **Data set reference (doi)** |
| --- | --- | --- | --- | --- | --- | --- | --- |
| Dytiscidae | Diving beetles | 38 | 3 | 2111 | M,N | Bergsten et al. (2013) | 10.5061/dryad.s631d |
| Dasypodidae | Armadillos | 13 | 5 | 6070 | M,N | Delsuc et al. (2003) | 10.5061/dyad.1838 |
| *Ensatina* | Salamanders | 69 | 2 | 823 | M | Devitt et al. (2013) | 10.5061/dryad.k9g50 |
| Muscidae | Flies | 39 | 3 | 1635 | M,N | Dsouli et al. (2011) | 10.5061/dryad.9025 |
| Chironomidae | Midges | 74 | 4 | 2701 | M,N | Ekrem et al. (2010) | 10.1016/j.ympev.2010.06.006 |
| Saxifragales | Part of core eudicots | 40 | 5 | 9005 | C,N | Fishbein et al. (2001) | 10.5061/dryad.684 |
| *Nothophagus* | Beeches | 51 | 6 | 5444 | C,N | Sauquet et al. (2012) | 10.5061/dryad.qq106tm4 |
| *Lycodon* | Wolf snakes | 61 | 3 | 2697 | M,N | Siler et al. (2013) | 10.5061/dryad.cp6gg |
| Neornithes | Modern birds | 161–200 | 255 | 361–2316 (mean = 1524, median = 1636) | N | Prum et al. (2015) | 10.5281/zenodo.28343 |
| Marsupialia | Marsupials | 35 | 1500 | 141–3660 (mean = 559.3, median = 429) | N | Duchêne et al. (2018) | 10.5061/dryad.353q5 |

M = mitochondrial; N = nuclear; C = chloroplast.

We also analysed two phylogenomic data sets that each comprised sequences from hundreds of loci (Table 2). The first data set consisted of sequences of a mixture of coding and non-coding regions from up to 200 bird species, representing all of the major extant lineages (Prum et al. 2015). The second data set comprised exon sequences from 35 marsupials, representing 18 of the 22 extant families (Duchêne et al. 2018). Each exon was further partitioned by codon position. We randomly split the phylogenomic data into alignments of 15 loci each to gain insight into the variation within them and for computational efficiency. The bird data and marsupial data were thus split into 17 and 300 smaller data sets, respectively.

We analysed each data set using maximum likelihood in IQ-TREE v1.6.7 (Nguyen et al. 2015), under each of the seven treatments described above (Table 1). The fit of the seven models was compared using the Bayesian information criterion (BIC). Under each treatment, we also examined estimates of evolutionary parameters, including the sum of the inferred branch lengths (tree length) and the proportional contribution of internal branches to the tree length (stemminess; Fiala and Sokal 1985). For analyses of each data set, we computed the path-distance metric between trees (Steel and Penny 1993) in a pairwise fashion across models of branch lengths. For the two phylogenomic data sets, we also compared each topological estimate with the maximum-likelihood estimate from the total data set, as reported in the original phylogenomic studies (Prum et al. 2015; Duchêne et al. 2018). We report comparisons across trees for each data set using multidimensional scaling of the pairwise distances between trees in two dimensions. The data sets, scripts used for analysis, and output files are available online (github.com/duchene/branch_length_models).

### Simulation Study

We conducted a simulation study to test for an association between the fit of different models of branch lengths and the length of sequences and number of taxa in the data set. As sequence length increases, there is more information available to identify the underlying evolutionary model. Similarly, an increasing number of taxa provides more information about the possible distribution of branch lengths, although a model with unlinked branch lengths across loci will gain large numbers of additional parameters. To explore the patterns of model support across these variables, we simulated sequence evolution along trees with varying numbers of taxa (4, 8, 16, and 32) and per-locus sequence length (500, 1000, 2000, and 4000 nucleotides). We started from symmetric time-trees with branch lengths of 10 million years (Myr). To convert these trees into phylograms, we multiplied the branch lengths (in time units) by branch rates drawn from a lognormal distribution using the R package NELSI (Ho et al. 2015). The scripts of the NELSI package are available online (github.com/sebastianduchene/NELSI), as are the scripts used for simulations and the output of our analyses (github.com/duchene/branch_length_models).

Using the framework described above, we simulated sequence evolution to produce pairs of loci under three different models of branch lengths. In the first model, the two gene trees had unlinked branch lengths but shared the same sum of branch lengths (tree length). Each set of branch rates was drawn from a lognormal distribution with mean 0.01 substitutions/site/Myr and log standard deviation of 0.2. In the second model, the two gene trees had proportionally linked branch lengths. This involved the two trees having the same pattern of branch-length variation but different tree lengths. The substitution rates of the two loci were 0.01 and 0.011 substitutions/site/Myr, without any rate variation across branches. In the third and final model, the two gene trees had unlinked branch lengths with different tree lengths. In this case, the two sets of branch rates were drawn from distributions with means of 0.01 and 0.011 substitutions/site/Myr, both with a log standard deviation of 0.2. This scenario is expected to be the most realistic representation of the evolutionary process. After the branch rates had been assigned, they were multiplied by the branch lengths of the time-trees. The resulting phylograms were used for our simulations of sequence evolution, which were performed using a Jukes-Cantor substitution model in the R package phangorn (Schliep 2011).

We generated 100 sets of branch rates and sequence alignments under each of the 48 combinations of branch-length model, number of taxa, and per-locus sequence length. The sequence alignments were then analysed using IQ-TREE. We used the BIC to compare the fit of three models of branch lengths, in which branch lengths were universally shared, proportionally linked, or unlinked across loci. In all cases, we assigned a separate substitution model to each locus. These scenarios correspond to treatments 3, 5, and 7 in our analyses of empirical data (Table 1). We also calculated the tree lengths and stemminess for the inferred trees and compared these with the metrics computed from the trees used for simulations of sequence evolution.

## Results

### Multilocus and Phylogenomic Data

In our analyses of multilocus and phylogenomic data sets, we found that the simplest model of universally shared branch lengths (treatment 1) provided a generally poorer fit than most other treatments (Fig. 2). As expected, this model also tended to have the lowest likelihood, and automatic model selection based on BIC rarely chose this model (Supplementary Fig. S1). For several data sets, including most of the multilocus data sets and the phylogenomic data set from birds, this model also led to longer terminal branches compared with the gene trees inferred using other models (Supplementary Fig. S1). In the case of some multilocus data sets, the simplest branch-length model also led to an estimate of the tree topology that was different from those obtained using the more complex models (Fig. 3a, 3f, and 3g).

**Figure 2.**
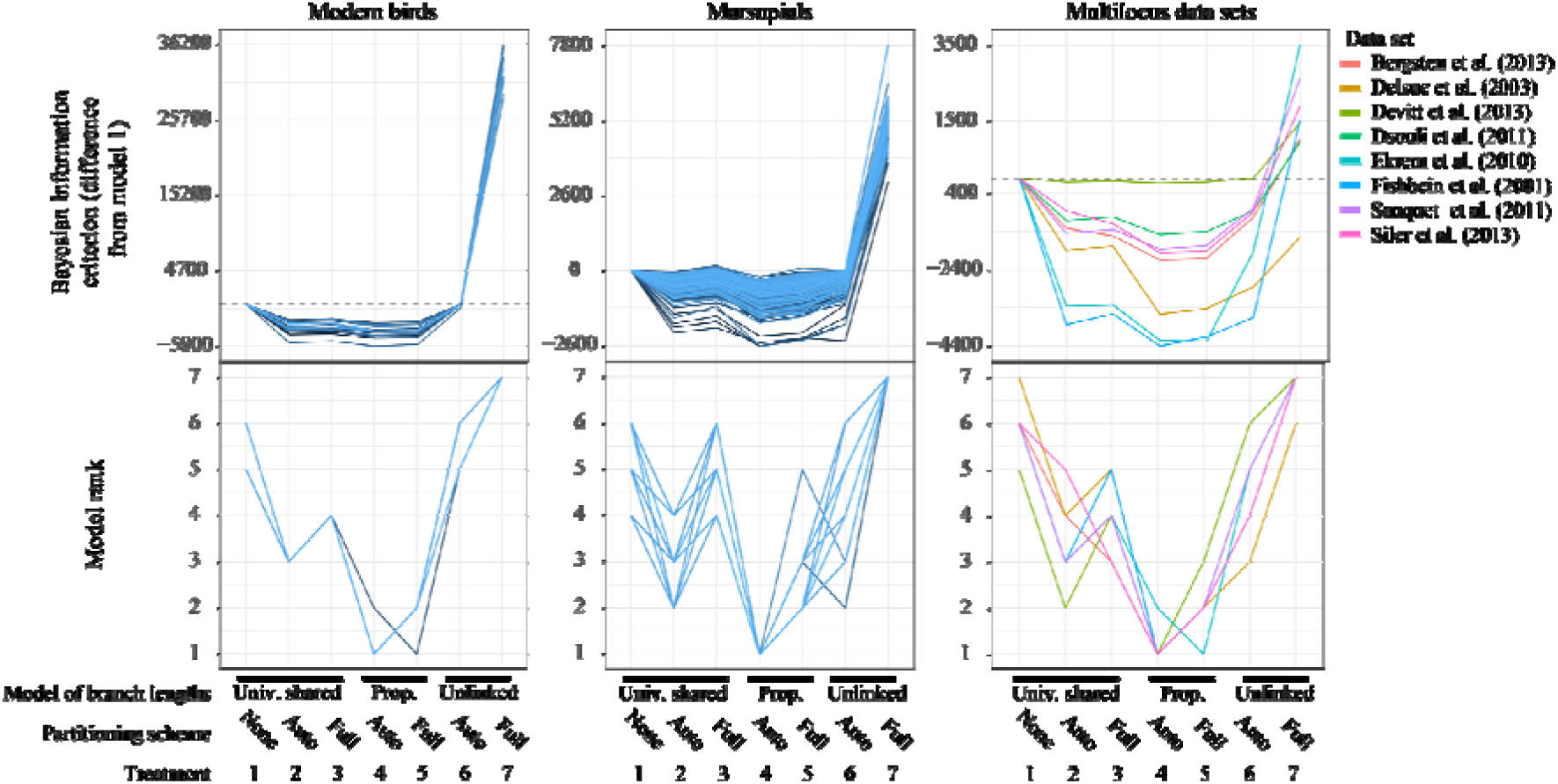
Statistical fit of seven models of nucleotide substitution and branch lengths across loci. The top row shows the relative statistical support for each treatment, measured in terms of the difference in the Bayesian information criterion (BIC) score from the simplest treatment (treatment 1). The bottom row shows the rank of each treatment in terms of its BIC score, with 1 representing the best-fitting treatment and 7 representing the worst-fitting treatment. Results are shown for analyses of eight multilocus and two phylogenomic data sets. The phylogenomic data comprise 17 data sets from birds and 300 data sets from marsupials. Each of these data sets comprises nucleotide sequences from 15 loci.

**Figure 3.**
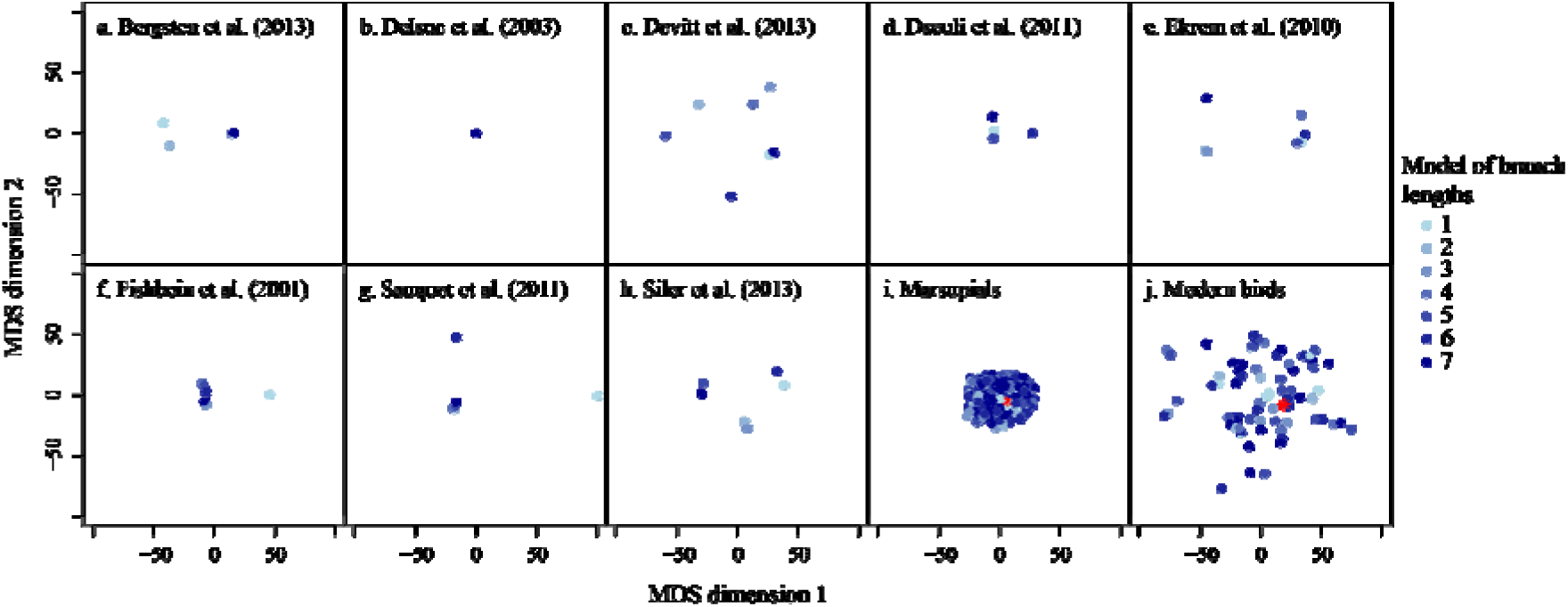
Two-dimensional representations of the topological path-distance between the trees inferred using each of the seven models of branch lengths. Distances between trees are represented after performing dimensionality reduction using multi-dimensional scaling (MDS). Red points in panels i and j indicate the maximum-likelihood estimates from the phylogenomic studies that first reported the data sets from marsupials (i) and modern birds (j).

A model with proportionally linked branch lengths (treatments 4 and 5) yielded the lowest BIC scores across the empirical data sets examined (Fig. 2). Specifically, the best-fitting model was the one in which branch lengths were proportionally linked and in which selection of the partitioning scheme was automated (treatment 4; Table 1). In addition to yielding the lowest BIC scores, the model with proportionally linked branch lengths tended to produce gene trees that were comparatively short, but with intermediate stemminess and levels of branch support (Supplementary Fig. S1). The second-best statistical fit was provided by a model in which branch lengths are shared across all loci, but where a separate substitution model is assigned to each locus.

The model with unlinked branch lengths across loci (treatment 7), which contained the largest number of parameters, consistently provided the poorest fit across all of the empirical data sets according to BIC scores (Fig. 2). Although this model had the highest likelihood (Supplementary Fig. S1), the penalty for its large number of parameters outweighed its improvement in likelihood. Nevertheless, this parameter-rich model did not lead to particularly distinct topological inferences, nor to greater distances from the reference bird and marsupial topologies when compared with the other models of branch lengths (Fig. 3).

The poor performance of the most complex model of branch lengths is also evidenced by the fact that automatic model selection often chose the simplest model (universally shared branch lengths). For the bird phylogenomic data, analyses using the most complex model consistently led to a greater contribution of internal branches to total tree length, lower mean bootstrap support across nodes, and a greater range in bootstrap support values across nodes (Supplementary Fig. S1).

### Simulation Study

In our analyses of sequence data generated by simulation, we found the expected pattern of an increasing preference for more parameter-rich models of branch lengths with increasing sequence length (Fig. 4). We also found that parameter-rich models were frequently selected when the data had increasing numbers of taxa. Regardless of the simulation conditions, a simple model with universally shared branch lengths was usually preferred when the sequences were very short (500 nucleotides) and when there were fewer than 32 taxa in the data set.

**Figure 4.**
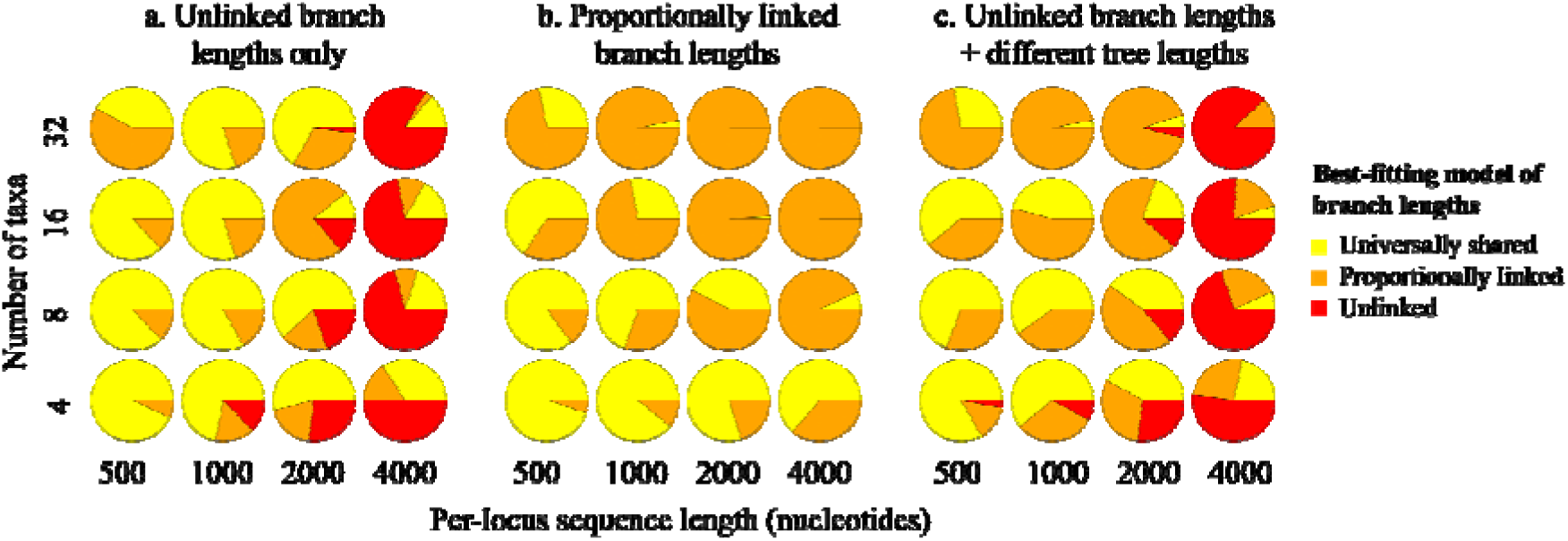
Comparison of branch-length models for two-locus data sets generated by simulation under three scenarios: (a) different patterns of branch lengths but identical tree lengths across gene trees; (b) identical patterns of branch lengths but different tree lengths across gene trees; and (c) different patterns of branch lengths and different tree lengths across gene trees. Each pie chart shows the proportion of 100 replicates for which each of the three models of branch lengths was selected using the Bayesian information criterion.

Under our first simulation scenario, in which loci had evolved with unlinked branch lengths but with the same tree length, the correct model of branch lengths was only preferred when each locus was 4000 nucleotides in length (Fig. 4a). In the second simulation scenario, in which the gene trees of the two loci had linked branch lengths with different tree lengths, the correct model of proportionally linked branch lengths was preferred when the number of taxa was greater than four (Fig. 4b). Finally, in the third simulation scenario, in which the two loci had gene trees with unlinked branch lengths and different tree lengths, the correct model with unlinked branch lengths was preferred when the loci were 4000 nucleotides in length (Fig. 4c). For shorter sequences and large numbers of taxa, a model with proportionally linked branch lengths was often chosen.

Across our simulation scenarios, we found branch-length estimates to be close to the true values (mean across loci), regardless of the model of branch lengths that was used for analysis (Supplementary Figs. S2–S3). For each scenario, the best-fitting model did not consistently lead to the most accurate estimates of branch lengths (Supplementary Figs. S4–S5). Nonetheless, analysing the data using a model with universally shared branch lengths almost always yielded shorter gene trees, which often had short internal branches compared with the trees inferred using other models of branch lengths (Supplementary Figs. S6–S7). In addition to highly accurate estimates of branch lengths, the tree topology was estimated correctly in every analysis. These outcomes are likely to reflect the fact that we explored a relatively narrow set of simulation parameters, despite this range being sufficient to produce variable impacts on model selection.

## Discussion

Our study has demonstrated that some degree of data partitioning is appropriate for improving model fit in phylogenetic analyses of multilocus data sets. In particular, our phylogenetic analyses of a range of empirical data sets showed that a model with proportionally linked branch lengths almost always provided the best fit. This outcome suggests that the dominant form of evolutionary rate variation that is being appropriately modelled is that across loci (i.e., gene effects), whereas the pattern of rate heterogeneity among branches does not vary enough across loci to warrant the use of a parameter-rich model with unlinked branch lengths. The model with proportionally linked branch lengths that was most often favoured in our analyses is available in several software packages (e.g., PhyML, Guindon et al. 2010; IQ-TREE, Nguyen et al. 2015), but not in others (RAxML, Stamatakis 2014).

Our results are broadly consistent with those of previous studies that identified biases in phylogenetic inference caused by underparameterization of the substitution model (Yang 1996; Lemmon and Moriarty 2004; Brandley et al. 2005; Revell et al. 2005; Marshall et al. 2006; Kainer and Lanfear 2015). Nonetheless, we have also found that unlinking branch lengths across loci incurs a substantial cost by introducing large numbers of parameters, leading to poor model fit. Unlinking branch lengths across loci led to estimates of topology and branch lengths with greater uncertainty than did models with intermediate numbers of branch-length parameters.

One way to identify an appropriate level of parameterization is to consider models of branch lengths with intermediate complexity to those considered here. For example, rather than estimating a separate, unlinked set of branch lengths for each locus, one might consider a model in which an intermediate number of groups of unlinked branch lengths are estimated. Each group of branch lengths can then be applied to multiple loci with a rate multiplier (i.e., proportional branch lengths) for each locus in the set. Some existing programs allow the specification of such intermediate models (e.g., PhyML Guindon et al. 2010). However, an algorithm to optimize the number of groups of unlinked branch lengths and their assignment to loci remains unavailable.

The results of our simulation study show that the most parameter-rich models are favoured only under certain conditions. Unlinking branch lengths across loci is an appropriate strategy only for data sets that comprise long sequences from moderate to large numbers of taxa (at least 32 taxa in our simulations). These large data sets contain the greatest amount of information about the distribution of rates across taxa. However, we would expect that a model with fully unlinked branch lengths would be strongly disfavoured for data sets with large numbers of loci, such as those encountered in phylogenomic studies.

Our study provides some insights into the importance of accounting for heterogeneity in molecular evolution across the genome. Variation in patterns of branch lengths across loci, as modelled in treatments 6 and 7 in our analyses, are the product of interactions between gene effects and lineage effects (Gillespie 1991; Cutler 2000; Gaut et al. 2011). Given that this description of rate variation across loci is perhaps the most biologically plausible, it is striking that the performance of this model is consistently poor across a wide range of multilocus data sets. One explanation for this result is that drivers of rate heterogeneity across lineages (e.g., differences in generation time) are largely independent of drivers of rate heterogeneity across loci (e.g., selective constraints). However, a more likely reason for the rejection of unlinked branch lengths is that such a model can involve enormous numbers of parameters, especially when the data set contains a large number of loci. As observed in our simulation study, this model is preferred only when each locus has a large number of nucleotide sites.

The findings of our study have implications for the use of clock models in molecular dating. Clock models describe the pattern of rate variation across the phylogeny, with relaxed clocks allowing a distinct rate along each branch (Ho and Duchêne 2014). When a separate relaxed-clock model is assigned to each locus, the number of parameters grows rapidly. Some studies have indicated that the careful assignment of a small number of clock models to subsets of the data can yield substantial improvements in model fit (e.g., Ho and Lanfear 2010; Duchêne and Ho 2014). However, the precision of divergence-time estimates is expected to improve with the number of loci (Zhu et al. 2015; Foster and Ho 2017; Angelis et al. 2018). Our results suggest that allowing different loci to share a single clock model is a reasonable approach, provided that the loci are allowed to have different relative rates. This approach is analogous to the model with proportionally linked branch lengths that has been considered here.

One of the assumptions in our analyses is that all of the loci have gene trees with identical topologies. This excludes the possibility of gene-tree discordance caused by incomplete lineage sorting, hybridization, or introgression. Discordance among gene trees leads to statistical inconsistency in phylogenetic analyses of concatenated data sets (Kubatko et al. 2007), and should be explicitly considered where possible (Mirarab et al. 2016). Forcing incongruent gene trees to share the same topology leads to distortions in the estimates of branch lengths (Mendes and Hahn 2016). Under these conditions, we might expect to see greater support for unlinking branch lengths across loci. The effect of variation in the topological signal across loci on models of branch lengths will require further investigation. Nonetheless, our results suggest that the variation in rates across loci and
lineages will often be well approximated by a model with proportionally linked branch lengths in analyses of concatenated sequence data.

## Conclusions

Our study has demonstrated the superior performance of phylogenetic models that proportionally link branch lengths across loci and that automate the process of selecting the data-partitioning scheme. Under- and overparameterization of the branch lengths across the gene trees can have negative impacts on phylogenetic analyses of multilocus data sets. For this reason, we recommend that proportionally linking branch lengths should be the default approach to analysing multilocus data sets. Our recommendations can be extended to phylogenomic data sets comprising large numbers of loci and taxa. Further examinations of the impact of branch-length models on divergence-time estimates, along with the effects of gene-tree discordance, are likely to be useful for improving the accuracy and precision of phylogenomic inferences.

## Acknowledgements

This work was supported by funding from the Australian Research Council to D.A.D. and S.Y.W.H. (grant DP160104173). S.D. was supported by a McKenzie Fellowship from the University of Melbourne. The authors acknowledge the Sydney Informatics Hub and the University of Sydney’s high performance computing cluster Artemis for providing the high-performance computing resources that have contributed to the research results reported in this paper.

